# Buffering gene expression noise by microRNA based feedforward regulation

**DOI:** 10.1101/310656

**Authors:** Pavol Bokes, Michal Hojcka, Abhyudai Singh

## Abstract

Cells use various regulatory motifs, including feedforward loops, to control the intrinsic noise that arises in gene expression at low copy numbers. Here we study one such system, which is broadly inspired by the interaction between an mRNA molecule and an antagonistic microRNA molecule encoded by the same gene. The two reaction species are synchronously produced, individually degraded, and the second species (microRNA) exerts an antagonistic pressure on the first species (mRNA). Using linear-noise approximation, we show that the noise in the first species, which we quantify by the Fano factor, is sub-Poissonian, and exhibits a nonmonotonic response both to the species lifetime ratio and to the strength of the antagonistic interaction. Additionally, we use the Chemical Reaction Network Theory to prove that the first species distribution is Poissonian if the first species is much more stable than the second. Finally, we identify a special parametric regime, supporting a broad range of behaviour, in which the distribution can be analytically described in terms of the confluent hypergeometric limit function. We verify our analysis against large-scale kinetic Monte Carlo simulations. Our results indicate that, subject to specific physiological constraints, optimal parameter values can be found within the mRNA-microRNA motif that can benefit the cell by lowering the gene-expression noise.

## 1 Introduction

Gene regulatory circuits encode diverse mechanisms to counter stochasticity arising from low-copy numbers of circuit constituents. Perhaps the most well-known example of this is negative feedback realized via gene autoregulation, where an expressed protein inhibits its own transcription/translation [1–12]. While such negative autoregulation is quite ubiquitous for *E. coli* transcription factors [13], it is surprisingly rare for eukaryotic transcription factors [14]. It is possible that the time delays associated with transporting the protein from the cytoplasm to nucleus compromise the noise buffering properties of negative feedback.

An alternative option is an incoherent feedforward loop that has been shown to be effective in maintaining a desired expression level in spite of changes in gene dosage [15], or upstream fluctuations in transcription factor levels [16–19]. Interestingly, increasing evidence shows that many eukaryotic genes are regulated by a specific feedforward architecture – the transcribed intronic regions of a gene that are removed during splicing are further processed to make a microRNA that targets the same gene’s mRNA [20]. This creates a strong coupling between the two species both in the sense of stoichiometry, and also timing of production events. We systematically study how such coupling in a feedforward loop alters noise in mRNA copy numbers, and identify parameter regimes which provide the most (and least) effective noise suppression.

Cellular regulatory circuits can be represented, up to a suitable level of detail, by systems of chemical kinetics. Unfortunately, exact characterisations of the copy-number distributions in a reaction system are often unavailable. Systems operating at a complex-balanced equilibrium are a notable exception in that they admit tractable product-form distributions [21–24]. Steady-state distributions have also been characterised in terms of generating functions in a growing collection of simple models that are not complex-balanced [25–30]. Such representations typically involve the use of special mathematical functions [31].

Approximative methods often provide a viable alternative in analysing a reaction system if exact results are unavailable or intractable. The linear-noise approximation (LNA) and moment-closure methods can reveal useful insights into the noise behaviour even for relatively complex reaction networks [32–38]. The quasi-steady state (QSS) approximation often leads to formulation of simplified models that can be more amenable to exact characterisation [39–43].

Here we apply these methodologies to analyse a reaction-kinetics model of a feedforward loop. Section 2 formally introduces the model and reviews some of its essential features as identified by an LNA analysis. Section 3 contains the derivation of the LNA results and cross-validates them with large-scale kinetic Monte Carlo simulations. Section 4 focuses on a special case in which the model admits a product-form distribution predicted by the Chemical Reaction Network Theory [23, 24]. Section 5 introduces a QSS approximation and shows that it can outperform the LNA in a specific parametric regime. The paper ends with a discussion of the current results and sketches lines of future inquiry.

## 2 The statement of the model and main results

We consider a discrete stochastic chemical kinetics system composed of two species X (mRNA) and Y (microRNA) which are subject to reaction channels

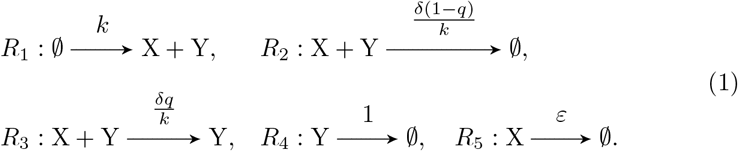

The molecules X and Y are produced synchronously with rate constant *k* through the reaction channel *R*_1_. In the reaction channels *R*_2_ and *R*_3_, a pair of X and Y molecules interact, which leads to the elimination of X; the Y molecule survives its antagonistic action on X with probability *q*, where 0 ≤ *q* ≤ 1 are allowed. For studies with a decoupled production of X and Y and *q* = 0 we refer the reader e.g. to [44, 45].

The strength of the antagonistic interaction is measured by the parameter *δ*. The reaction rates of both second-order reactions are divided by *k* in order to achieve a classical scaling [46] of the reaction system (1) with respect to the parameter *k*. Doing so makes the ensuing analysis more transparent.

The reaction channels *R*_4_ and *R*_5_ describe the spontaneous degradation of Y and X. By measuring time in units of the expected lifetime of Y, we are able to fix the reaction constant of *R*_4_ to one. The parameter *ε* gives the ratio of Y to X lifetimes. In particular, small values of *ε* pertain to the assumption that X be much more stable than Y.

In this paper we aim to examine how the choice of parameter values affects the stochastic noise in the X copy number at steady state. We use the Fano factor, which is defined as the ratio of the variance to the mean, as our chosen noise metric. It is well known that the Fano factor is equal to one for the Poisson distribution. We refer to a distribution with Fano factor lower than one as sub-Poissonian.

Using the linear-noise approximation (LNA), we obtain for the Fano factor

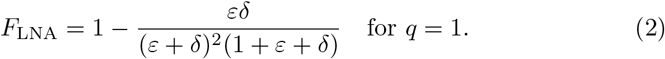

The details of the derivation follow in Section 3. Note that the LNA result (2) is independent of the parameter *k*. The function (2) is visualised as a heat map in the upper-left panel of Fig. 1.

**Fig. 1.**
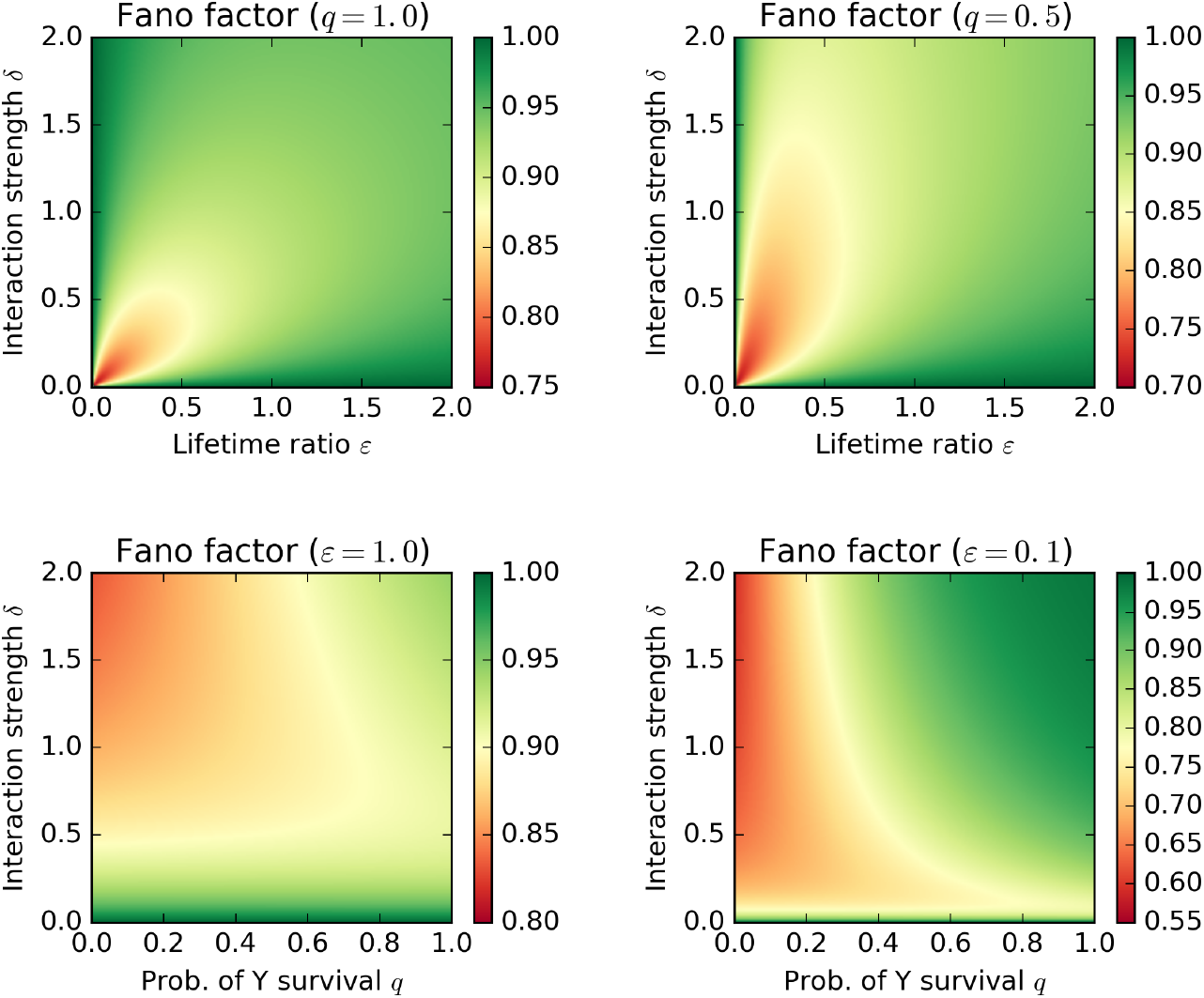
The Fano factor of species X (mRNA) by linear-noise approximation (LNA) depending on the model parameters.

Elementary analysis of (2) reveals that

1. The steady-state distribution of X is sub-Poissonian if *ε* > 0 and *δ* > 0.
2. If there is no interaction (*δ* = 0) or if X is stable (*ε* = 0), the Fano factor is equal to one.
3. For a fixed positive value of *ε*, the Fano factor is a nonmonotonic function of *δ*, initially decreasing before reaching a minimum, and slowly increasing back to one as *δ* → ∞.
4. The analogous holds if the roles of *ε* and *δ* are reversed in Property 3.
5. The function *F*_LNA_(*ε, δ*) is discontinuous at (*ε, δ*) = (0, 0). A range of limiting values can be achieved depending on along which ray the origin is approached.
6. The function *F*_LNA_(*ε, δ*) does not have an unconstrained minimum. An infimum of 0.75 is approached if *ε* = *δ* → 0.

The function *F*_LNA_ can be efficiently evaluated also for 0 ≤ *q* ≤ 1, but the algebra reveals little. Graphical examination of *F*_LNA_ indicates that all the above properties hold for 0 ≤ *q* ≤ 1 (see the upper-right panel of Fig. 1 for *q* = 0.5). The infimum of the Fano factor in Property 6 is lower than 0.75 if *q* < 1 and is approached along a different ray emanating from the origin. In the exceptional case *q* = 0 the dependence of *F*_LNA_ on *ε* and *δ* is monotonous (the lower panels of Fig. 1).

Property 2 suggests, but does not provide a definitive proof, that the distribution of X is Poissonian if *ε* = 0 or *δ* = 0. In the non-interaction case (*δ* = 0), the proof is straightforward: the dynamics of X is that of a simple immigration- and-death process, which is known to generate a Poisson distribution [47–49]. If X is stable (*ε* = 0), the distribution is again Poisson, but the proof requires a more subtle reasoning based on the Chemical Reaction Network Theory. We present the details in Section 4.

The discontinuity of *F*_LNA_(*ε, δ*) at the origin indicates that a surprisingly rich behaviour, in terms of the chosen noise metric, can be recovered by focusing solely on the *ε, δ* ≪ 1 parametric region. We pursue this line of inquiry in Section 5, where we identify a family of discrete distributions, which describe the limit behaviour of (1) in this parametric region. For reasons made explicit later, we refer to this description as the quasi-steady-state (QSS) model. We demonstrate that the QSS model can be superior to the LNA in predicting simulation results.

## 3 Linear-noise approximation

In linear-noise approximation, the mean behaviour is given by the law-of-mass-action formulation of the reaction system (1), which is

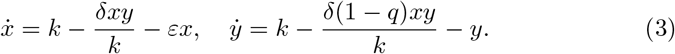

Setting the derivatives in (3) to zero and solving the resulting algebraic system in *x* and *y* yield the stationary mean values

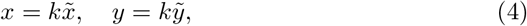

where

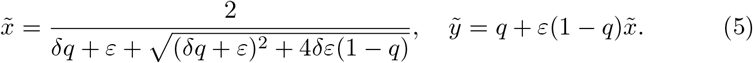

Note that the means (4) scale with the production rate constant *k*.

In order to obtain the LNA of the variance, we need to determine the steady-state fluctuation and dissipation matrices of the reaction system (1). The dissipation matrix ***A*** is equal to the linearisation matrix of the system (3), i.e.

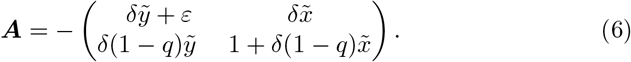

The fluctuation matrix ***B*** is obtained in the following manner [50]: for each reaction channel in the system (1), we calculate the outer product of the reaction vector [51, 52] with itself, and multiply it by the (steady-state) reaction rate; then we sum the results over all reaction channels. In our particular example this leads to

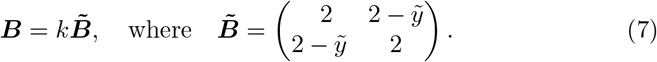

The fluctuation–dissipation theorem [53] states that the covariance matrix ***Σ*** of the random vector of steady-state X and Y copy numbers satisfies

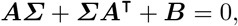

i.e

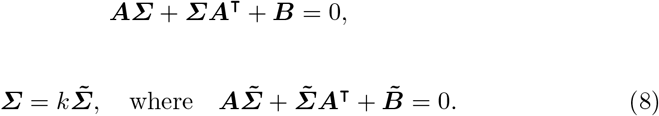

Note that the covariance matrix (8) scales with the production rate constant *k*. The linear algebraic system (8) can be written in a flattened form as

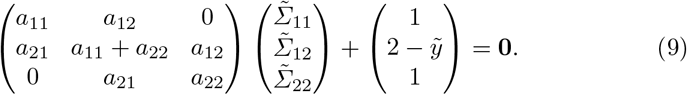

The Fano factor is given by

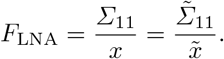

Since both the mean *x* and the variance *Σ*_11_ scale linearly with *k*, the Fano factor is independent of the parameter.

For 0 ≤ *q* ≤ 1, we solve (9) for every combination of parametric values numerically by a fast linear-algebra solver. For *q* =1, equations (5) and (6) simplify to

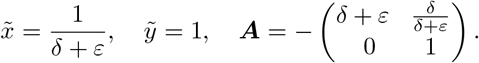

The linear system (9) becomes upper triangular, and the formula (2) is obtained after few elementary steps.

A classical system-size-expansion argument [54] guarantees that the LNA accurately describes the reaction system (1) as *k* tends to infinity. We demonstrate the asymptotics in Fig. 2, in which we compare the LNA of the Fano factor to the value obtained by the application of Gillespie’s stochastic simulation algorithm (SSA) [55]. We observe a perfect agreement between the two if *k* = 30 (Fig. 2, left panel). Although the agreement remains satisfactory for *k* = 3, the SSA results are now seen to deviate systematically from the LNA prediction for moderate values of *ε* and *δ* (Fig. 2, right panel).

**Fig. 2.**
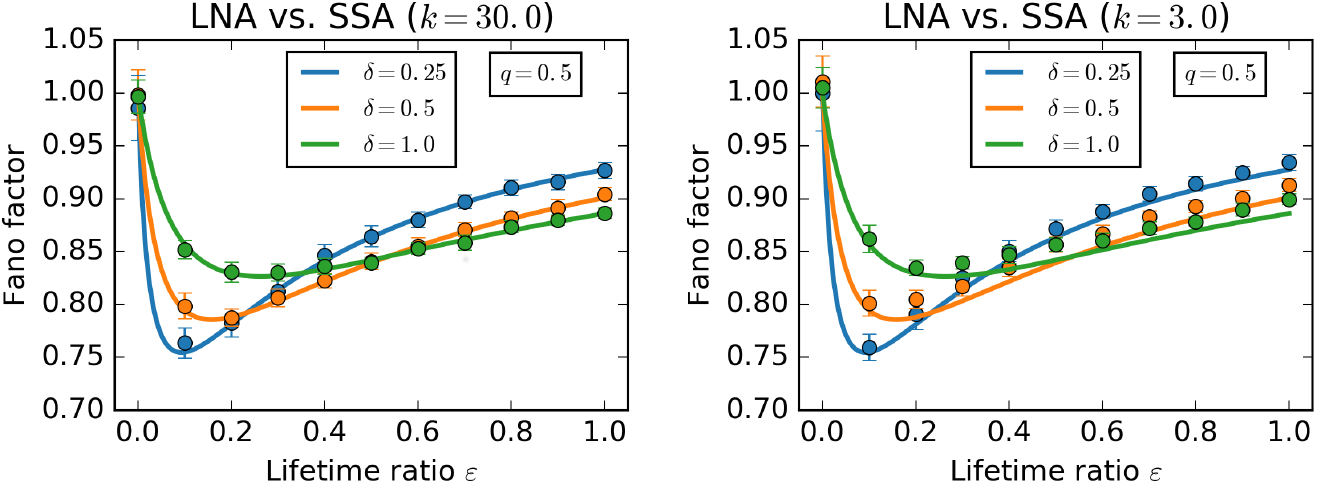
The Fano factor of species X (mRNA) as function of model parameters as given by the linear-noise approximation (LNA, solid lines) and stochastic simulation algorithm (SSA, discrete markers). Two values of *k* are used, one large (*k* = 30, left panel) and one moderate (*k* = 3, right panel). The LNA is independent of *k*. Error bars indicate 99.9% confidence intervals for the simulation-based Fano factors.

The reaction species mean values and standard deviations were calculated using Stochpy’s [56] implementation of Gillespie’s direct method [55]. We skipped over the first 30 units of time to avoid the influence of an initial transient; we estimated the moments from the next 10^5^ iterations (for *k* = 3) or 10^6^ iterations (for *k* = 30) of the algorithm. The Fano factor was calculated as the ratio of the squared standard deviation to the mean value. The procedure was repeated to obtain 25 independent Fano factor estimates. The repetition increased accuracy and facilitated the construction of confidence intervals.

## 4 Stability of X implies Poisson distribution

In Section 3, we reported that if Y has a chance of surviving the antagonism with X (*q* > 0) and if X is stable (*ε* = 0), then the LNA of the Fano factor is equal to one. Here we expand on this observation by providing an actual proof that the steady-state copy number of X (and also that of Y) follows the Poisson distribution. The argument is based on the application of the Chemical Reaction Network Theory (CRNT) [23].

Setting *ε* = 0 in the reaction system (1) is tantamount to removing the reaction channel *R*_5_ for spontaneous degradation of X. The four remaining reaction channels involve *N* = 3 complexes, namely the empty set 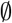, the pair X +Y, and the singleton Y. The three complexes are represented as vertices of the reaction graph (Fig. 3). The reaction graph has a single linkage class (*l* = 1). Given that the linkage class is strongly connected, the chemical system is weakly reversible in the sense of the CRNT.

**Fig. 3.**
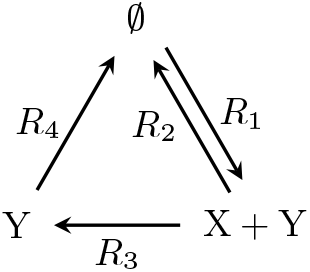
Graphical representation of a reduced reaction system obtained from (1) by disabling the spontaneous degradation of X (reaction channel *R*_5_). The reduced reaction system is seen to be weakly reversible (unless *q* = 0). Additionally, it has zero deficiency and admits no conservation laws. Therefore, the copy numbers of X and Y are independent and Poissonian.

The reaction vectors span the entire two-dimensional (*s* = 2) space of X and Y copy number pairs. The deficiency of the reaction network is obtained by the well-known formula *δ* = *N* – *l* – *s* = 0. According to CRNT [23], weakly reversible networks of zero deficiency admit a product-form steady-state distribution

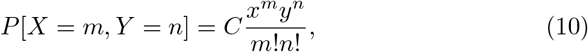

where *x* = *k*/*δq* and *y* = *kq* are obtained by solving the law-of-mass action kinetics (3) at steady state, i.e. by setting *ε* = 0 in (4)–(5).

Since the reaction system does not admit any conservation laws, the support of the distribution (10) includes all pairs of nonnegative integers *m,n* ≥ 0. The normalisation constant *C* is then readily determined as *C* = *e*^−*x*−*y*^. In other words, the joint distribution of X and Y is the product of the marginal distributions, either of which is Poissonian with mean *x* and *y*, respectively.

One of the powerful aspects of the Chemical Reaction Network Theory is that it generalises to complex extensions of Fig. 3 as long as they do not violate its fundamental structural properties. Biologically, microRNAs tend to be promiscuous binders [57]. In Appendix A we extend the network of Fig. 3 by an interaction between Y (the microRNA) and decoy binding sites. We show that the copy-number distributions remain Poissonian.

## 5 Quasi-steady-state approximation

The LNA results presented in Section 2 suggested that the reaction system (1) exhibits a wide range of noise behaviour in the small *ε, δ* ≪ 1 region. In this section we develop an approximative description to the reaction system (1) that is applicable for such parametric choices. Specifically, we assume that

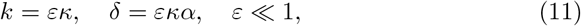

where *κ* and *α* are the rescaled production and interaction rate constants. The scaling (11) guarantees than *ε* and *δ* are both small and that they approach the origin along a ray with slope given by *κα*. Note that taking *ε* and *δ* to zero whilst keeping the production rate *k* fixed would have led to a divergence in the level of X. By making the production also scale with *ε* we are able to approach a distinguished limiting distribution as *ε* tends to zero. Also note that (11) removes the classical scaling with respect to the production rate constant from the system (1). It turns out that this choice keeps the ensuing analysis more tractable.

The steady-state probability distribution *p_m,n_* of observing *m* molecules of X and *n* molecules of Y in the system satisfies the master equation

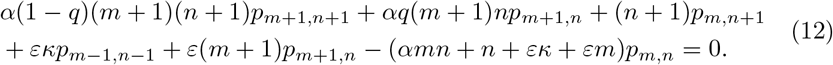

The first three terms in (12) give the probability influx into the reference state (*m, n*) due to the reaction channels *R*_2_, *R*_3_ (interactions) and *R*_4_ (decay of Y). These terms are not multiplied by *ε*, i.e. the three reactions are considered to be fast. The next two terms give the probability influx due to the channels *R*_1_ (production) and *R*_5_ (decay of X). These terms are of order *ε*, i.e. the two reactions are slow. The final, negative, term in (12) gives the probability efflux from the reference state (*m, n*). The master equation states that the influx and the efflux balance out.

The separation between fast and slow reactions suggests that we should use quasi-steady-state (QSS) reduction techniques to study (12). The molecule Y plays the role of the transient, highly reactive, species (or the QSS species [58, 59]). Each time a molecule of Y is produced, a brief period of interaction with X ensues, with ends with the elimination of Y (whether through natural degradation or through the interaction with X if *q* < 1).

We seek a power-series solution

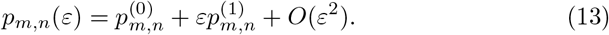

Inserting (13) into (12) and collecting *O*(1) terms yields

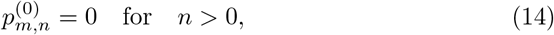

which states that the probability of observing a non-zero number of the QSS species Y is *O*(*ε*) small. Further analysis helps establish additional relations

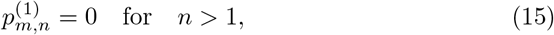

which state that the probability of observing two or more molecules of Y is *O*(*ε*^2^) small. Although relations (14) and (15) are sufficient for our present analysis, they can be generalised to 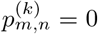 for *n* > *k*, which state that the probability of having more than *k* molecules of Y is *O*(*ε*^*k*+1^).

In light of (14), it remains to determine the terms 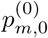 in order to obtain the limiting probability distribution. To this end, it is sufficient to use the master equation (12) for *n* = 0 and *n* =1, i.e.

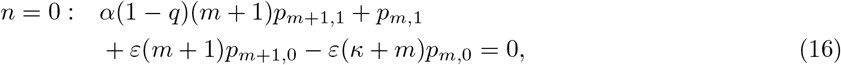

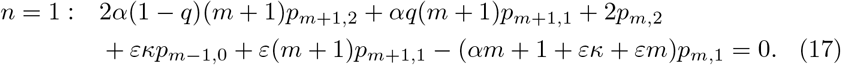

Inserting the power-series ansatz (13) into (16)–(17) and collecting *O*(*ε*) terms yields

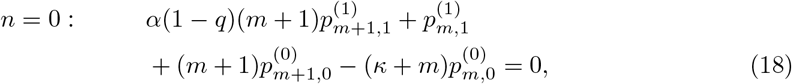

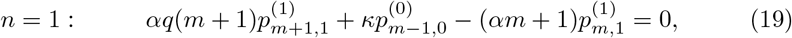

whereby we made use of the relations (14) and (15). In particular, the relations (15) guarantee that (18)–(19) forms a closed system of difference equations in the unknown series 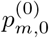 and 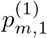.

In order to solve (18)–(19), we introduce the generating functions

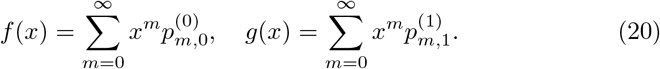

Multiplying (18)–(19) by *x^m^* and summing over *m* ≥ 0 yield a system of ordinary differential equations

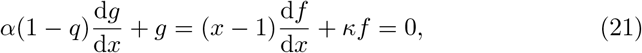

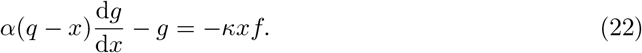

Next, we turn the system (21)–(22) of two first-order ordinary differential equations into a single second-order ordinary differential equation.

First, we eliminate *g* by adding the equations (21) and (22) up, and dividing the result by 1 – *x*, which gives

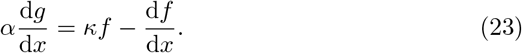

Second, we eliminate d*g*/d*x* by adding up the (*x* – *q*)-multiple of (21) and the (1 – *q*)-multiple of (22) before dividing the result by *x* – 1, whereby we obtain

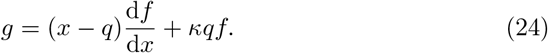

Differentiating (24), we find

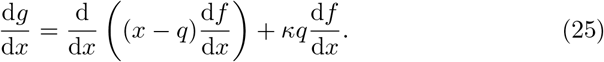

Combining (23) and (25), we arrive at an ordinary differential equation of the second order for *f*, which reads

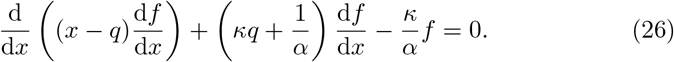

We look for a solution to (26) in the form of a power series

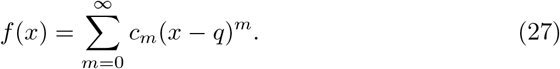

Inserting (27) into (26), we have

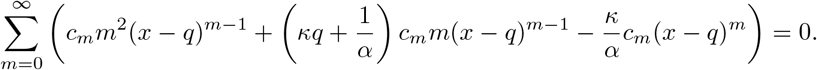

Equating like powers of (*x* – *q*) yields a recursion

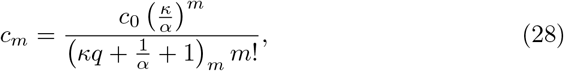

solving which yields

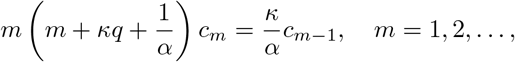

where

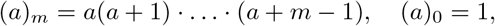

represents the *m*-th rising factorial from a number *a* > 0.

Substituting (28) into (27) we find that

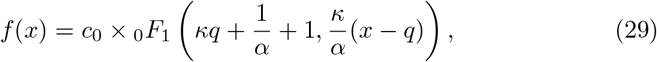

in which the confluent hypergeometric limit function _0_*F*_1_ is defined by the convergent series

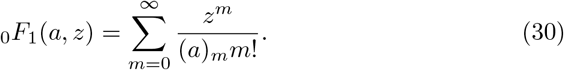

Imposing the normalisation condition *f*(1) = 1, we determine the prefactor *c*_0_ in (29) and obtain

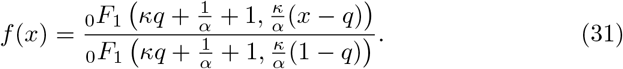

Basic properties of the confluent hypergeometric limit function can be established using its power-series representation (30). Repeatedly differentiating the series (30) term by term yields

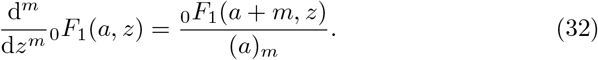

Comparing (30) with the power-series expansions of the normal and modified Bessel functions [31], we obtain

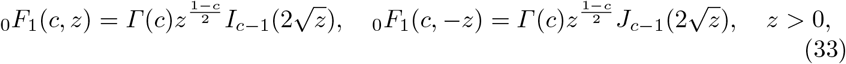

where *Γ*(*z*) is the gamma function, *J_ν_*(*z*) is the Bessel function, and *I_ν_*(*z*) is the modified Bessel function of order *ν*.

Repeatedly differentiating (31) yields

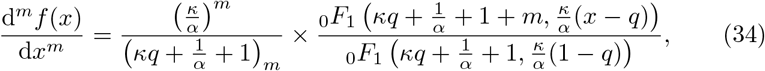

which provides an approximation

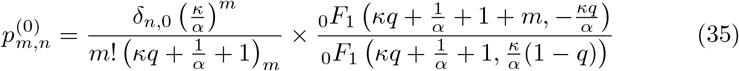

for the desired solution to the master equation (12). Evaluating the derivatives of *f*(*x*) at *x* = 1, we obtain the factorial moments [60]

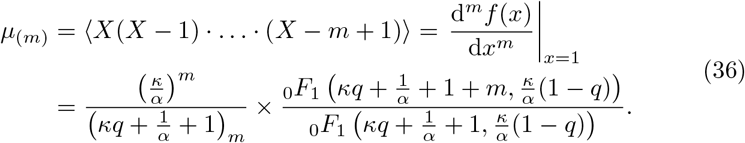

At the same time as noting that the mean 〈*X*〉 trivially coincides with the first factorial moment *μ*_(1)_, we also point out that the main characteristic of interest here, the Fano factor, can be expressed in terms of the first two factorial moments as

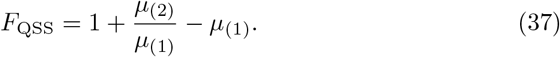

We expressly mention, without carrying out the somewhat tedious calculation, that the probability distribution (35) and the moments (36) can be written in terms of Bessel’s functions via (33).

In Fig. 4, we compare the values of the Fano factor obtained in the parametric regime (11) by the quasi-steady state (QSS) approximation, the linear-noise approximation (LNA), and by the application of stochastic simulation algorithm (SSA). We observe that the QSS model predicts the SSA results more faithfully than the LNA. For *q* > 0, the LNA overestimates the dip in the Fano factor (Fig. 4, left panel). For *q* = 0, the LNA predicts a monotonous decrease of the Fano factor with the interaction strength, whereas the QSS and SSA results both show an eventual slow increase (Fig. 4, right panel).

**Fig. 4.**
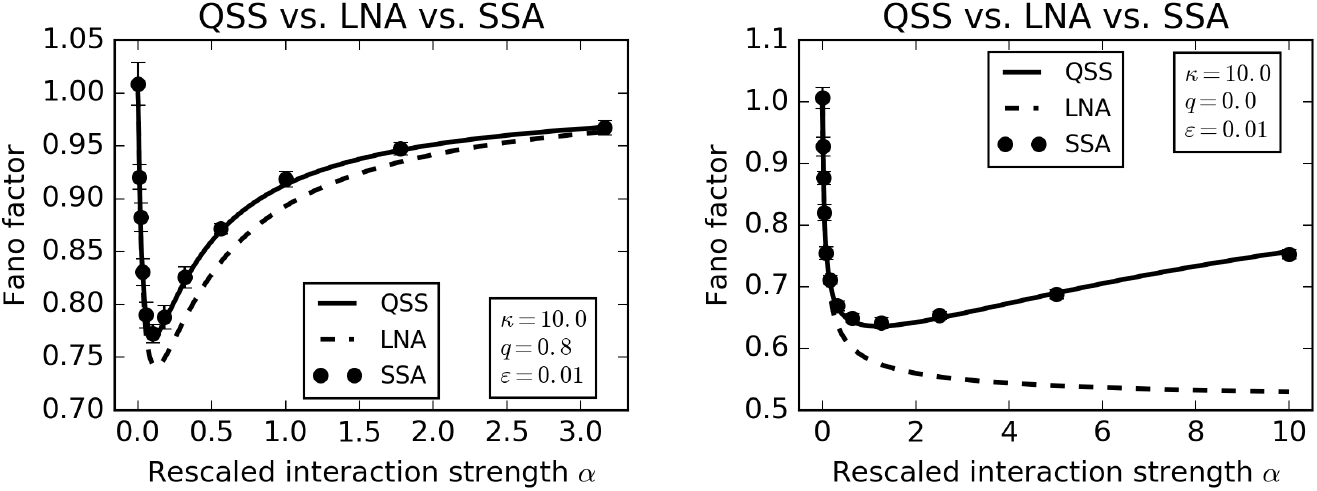
The Fano factor of species X (mRNA) as function of rescaled model parameters as given by the quasi-steady state (QSS) model, the linear-noise approximation (LNA) and the stochastic simulation algorithm (SSA). Error bars indicate 99.9% confidence intervals for the simulation-based Fano factors.

The LNA values were calculated by the method of Section 3, whereby the original parameters *k* and *δ* were recovered from the values of *κ* and *α* through relations (11). Each SSA value was computed in StochPy [56] from 25 independent sample paths each consisting of 10^5^ iterations of Gillespie’s direct method. The QSS values were obtained from (36)–(37).

In Fig. 5, we compare simulation-based estimates of the X copy-number distribution to the QSS approximation (35) and a Poissonian benchmark. The agreement between the simulation and the QSS results improves as the value of *ε* is decreased: compare the left panels with *ε* = 0.1 to the right panels with *ε* = 0.01. Consistently with our previous reports of the model’s sub-Poissonian behaviour, the species X copy number distributions are narrower than the Poissonian benchmark.

**Fig. 5.**
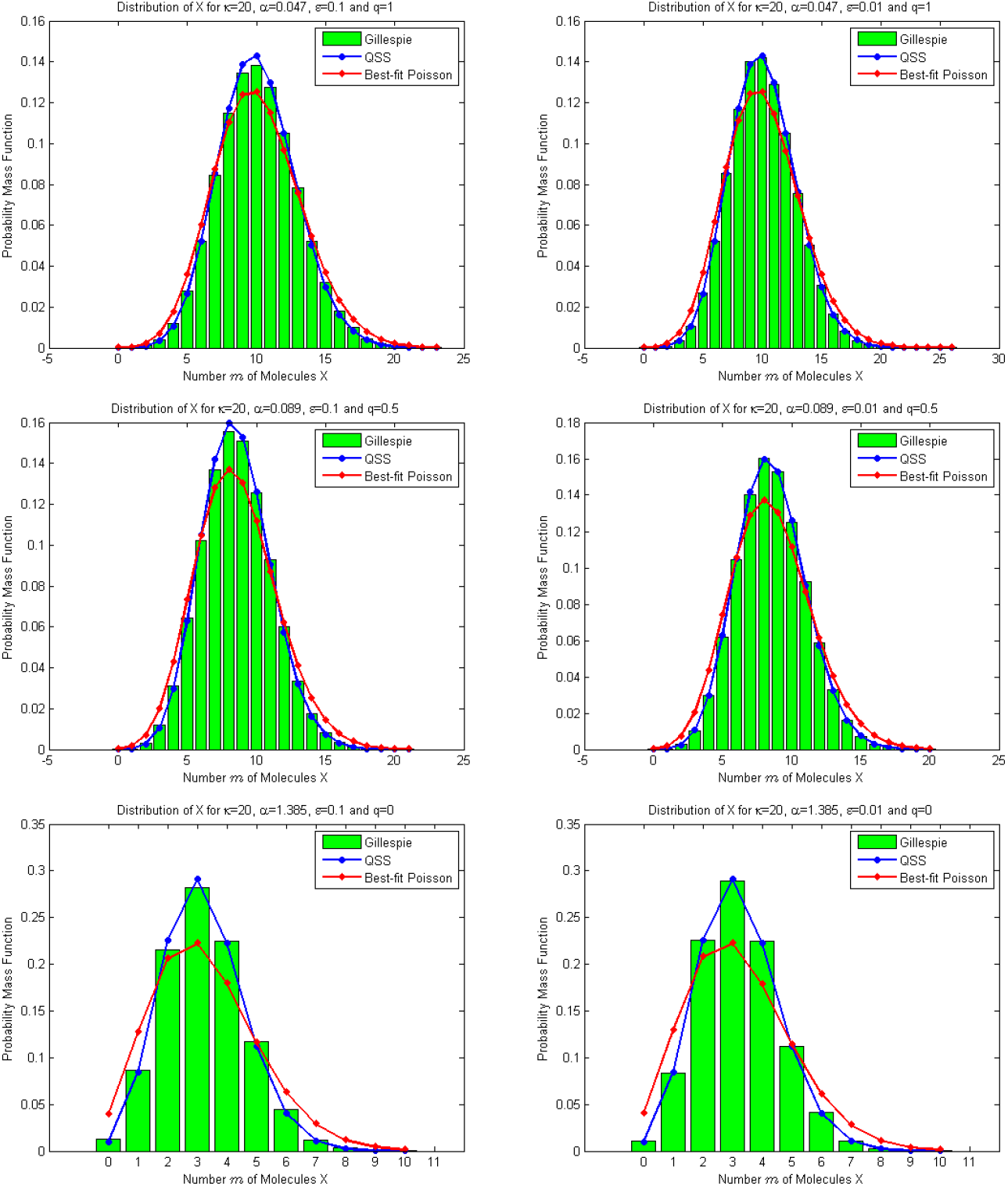
Marginal distributions of X copy number obtained by Gillespie’s algorithm, quasi-steady-state (QSS) approximation, and by maximum-likelihood fitting of a Poisson distribution.

The values of the interaction *α* were selected in Fig. 5 so as to minimise the Fano factor for the given values of *κ* and *q*. Each histogram is based on 10^5^ independent sample paths of the chemical system (1) generated with Gillespie’s direct method. The Poisson distribution was fit to the simulation data by maximum likelihood estimation.

## 6 Discussion

In this paper we considered a stochastic model for a feedforward loop driven by the interaction between mRNA and microRNA species. Using a combination of mathematical and computational methods, we investigated the effects of microRNA based regulation on the mRNA noise levels. The model behaviour depends on several parameters, namely: the gene-expression rate, the interaction strength, the microRNA to mRNA lifetime ratio, and the probability of microRNA surviving its interaction with mRNA.

Our results indicate that feedforward regulation can buffer mRNA noise to sub-Poissonian levels. The Fano factor (the variance to mean ratio) exhibits a nonmonotonic behaviour: for a fixed microRNA to mRNA lifetime ratio, there is an optimal value of the interaction strength that minimises the Fano factor; conversely, for a fixed interaction strength, there exists an optimal lifetime ratio. However, an unconstrained minimum with respect to the two parameters does not exist. The infimum can be approached by taking small microRNA to mRNA lifetime ratios and interaction strengths. Intriguingly, if mRNA is assumed to be completely stable, the Fano factor is equal to the Poissonian value of one. Decreasing the probability of microRNA survival in its interaction with mRNA leads to lower values of the Fano factor.

Much of the model behaviour has been identified using the linear-noise approximation. We also used additional methodologies to examine some of the phenomena more closely. Specifically, we used the Chemical Reaction Network Theory to prove conclusively that the mRNA copy-number distribution is Poisson if mRNAs are stable. Additionally, we constructed a quasi-steady-state (QSS) approximation of the model, which applies specifically to the situation of large mRNA to microRNA lifetimes and low interaction strengths. The QSS approximation was shown to outperform the LNA in this regime.

We do not consider the current model to be an exhaustive description of a microRNA based regulation of gene expression. Instead, our intention was to examine, using a minimalistic chemical system, the effects on the underlying feedforward regulation on the noise in the regulated species. In order to obtain more realistic and/or general formulations, we propose to extend the model in several specific directions.

First, we propose to extend the model by transcriptional bursting to investigate the effects of feedforward regulation on the super-Poissonian mRNA noise. Second, we propose to include translation, and examine the effects on protein noise. Analyses of different systems suggest that protein noise is not simply proportional to the mRNA noise, but also depends on mRNA autocorrelation times [61]; proteins can also control their noise through transcriptional and/or translational feedbacks. Our third proposition is to consider the effect of non-specific binding [62] of microRNAs on their ability to regulate gene-expression noise. We have made a step in this direction in Appendix A, showing that the Poissonian case remains Poissonian after the inclusion of decoy binding sites. In more general sitations, the addition of non-specific binding is expected to lead to nontrivial effects, the understanding of which may require employing additional techniques [63–65].

In summary, we studied a stochastic feedforward loop featuring a coupled production and antagonistic interaction. We examined the consequences of the interaction on gene-expression noise. Using a combination of different methodologies we characterised the model behaviour in several parameter regions of interest. We expect that analogous approaches will be helpful to understand more complex versions of the model as well as other examples of gene-regulatory motifs operating at low copy numbers.

## Appendix A. Decoy binding sites

The Chemical Reaction Network Theory (CRNT) can be applied to reaction networks that extend Fig. 3 with additional reaction channels as long as they do not violate its fundamental structural properties. Assume, for example, that a molecule Y can bind to a free binding site B to form a heterodimer C, and that the heterodimer can dissociate into its constituents Y and B. Consider a reaction network obtained by extending that of Fig. 3 by the reversible pair of reactions Y + B ⇄ C. It is clear that the extended network remains weakly reversible, and that it involves *N* = 5 different complexes (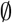, X + Y, Y, Y + B, and C), which are interconnected within *l* = 2 linkage classes. The copy numbers (*m, n, i, j*) of the four reaction species (X, Y, C, and B) are constrained by the conservation law *i* + *j* = *ν*, where *ν* gives the total number of binding sites. Consequently, the stoichiometric subspace has dimension *s* = 4 – 1 = 3, and the deficiency of the extended network is equal to *δ* = *N* – *l* – *s* = 5 – 3 – 2 = 0. It follows that the steady-state distribution has the product form [23]

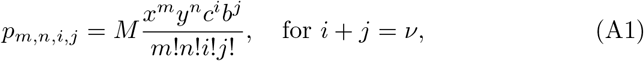

where

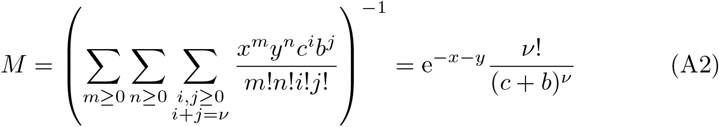

is the normalisation constant. The first two components of the complex-balanced equilibrium (*x, y, c, b*) are still given by the values *x* = *k*/*δq* and *y* = *kq* obtained in Section 4 in the absence of decoys. For the other two components of the complex-balanced equilibrium we have the balance condition *yb* = *Lc*, where *L* is the dissociation constant, and the conservation law *b* + *c* = *ν*; solving in *c* and *b* yields

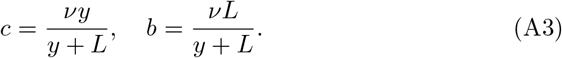

Inserting (A3) into (A1)–(A2) and simplifying yields

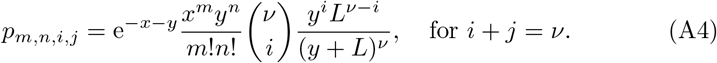

The distribution in (A4) is a product of two Poissons for X and Y copy numbers and of a binomial for the number of heterodimers C. In particular, the distribution of X (mRNA) remains Poissonian despite the addition of an unspecific interaction with decoy binding sites.

## Bibliography

[1] Voliotis, M., Bowsher, C.G.: The magnitude and colour of noise in genetic negative feedback systems. Nucleic Acids Res. (2012)

[2] Singh, A., Hespanha, J.P.: Optimal feedback strength for noise suppression in autoregulatory gene networks. Biophys. J. 96 (2009) 4013–4023

[3] Dublanche, Y., Michalodimitrakis, K., Kummerer, N., Foglierini, M., Serrano, L.: Noise in transcription negative feedback loops: simulation and experimental analysis. Mol. Syst. Biol. 2 (2006) 41

[4] Nevozhay, D., Adams, R.M., Murphy, K.F., Josic, K., Balazsi, G.: Negative autoregulation linearizes the dose response and suppresses the heterogeneity of gene expression. P. Natl. Acad. Sci. USA 106 (2009) 5123–5128

[5] Bokes, P., Singh, A.: Gene expression noise is affected differentially by feedback in burst frequency and burst size. J. Math. Biol. 74 (2017) 1483–1509

[6] Bokes, P., Lin, Y., Singh, A.: High cooperativity in negative feedback can amplify noisy gene expression. B. Math. Biol. (2018) doi: 10.1007/s11538-018-0438-y

[7] Rosenfeld, N., Elowitz, M.B., Alon, U.: Negative autoregulation speeds the response times of transcription networks. J. Mol. Biol. 323 (2002) 785–793

[8] Simpson, M.L., Cox, C.D., Sayler, G.S.: Frequency domain analysis of noise in autoregulated gene circuits. P. Natl. Acad. Sci. USA 100 (2003) 4551–4556

[9] Becskei, A., Serrano, L.: Engineering stability in gene networks by autoregulation. Nature 405 (2000) 590–593

[10] Swain, P.S.: Efficient attenuation of stochasticity in gene expression through post-transcriptional control. J. Mol. Biol. 344 (2004) 965–976

[11] Singh, A.: Negative feedback through mRNA provides the best control of gene-expression noise. IEEE T. NanoBiosci. 10 (2011) 194–200

[12] Thattai, M., van Oudenaarden, A.: Intrinsic noise in gene regulatory networks. P. Natl. Acad. Sci. USA 98 (2001) 151588598

[13] Alon, U.: Network motifs: theory and experimental approaches. Nat. Rev. Genet. 8 (2007) 450–461

[14] Stewart, A.J., Seymour, R.M., Pomiankowski, A., Reuter, M.: Underdominance constrains the evolution of negative autoregulation in diploids. Plos Comput. Biol. 9 (2013) e1002992

[15] Bleris, L., Xie, Z., Glass, D., Adadey, A., Sontag, E., Benenson, Y.: Synthetic incoherent feedforward circuits show adaptation to the amount of their genetic template. Mol. Syst. Biol. 7 (2011) 519

[16] Osella, M., Bosia, C., Corá, D., Caselle, M.: The role of incoherent microRNA-mediated feedforward loops in noise buffering. Plos Comput. Biol. 7 (2011) e1001101

[17] Tsang, J., Zhu, J., van Oudenaarden, A.: MicroRNA-mediated feedback and feedforward loops are recurrent network motifs in mammals. Mol. Cell 26 (2007) 753–767

[18] Strovas, T.J., Rosenberg, A.B., Kuypers, B.E., Muscat, R.A., Seelig, G.: MicroRNA-based single-gene circuits buffer protein synthesis rates against perturbations. ACS Synth. Biol. 3 (2014) 324–331

[19] Soltani, M., Platini, T., Singh, A.: Stochastic analysis of an incoherent feedforward genetic motif. In: 2016 American Control Conference (ACC). (2016) 406–411

[20] Bosia, C., Osella, M., Baroudi, M.E., Cora, D., Caselle, M.: Gene autoregulation via intronic microRNAs and its functions. Bmc Syst. Biol. 6 (2012) 131

[21] Jahnke, T., Huisinga, W.: Solving the chemical master equation for monomolecular reaction systems analytically. Journal of mathematical biology 54 (2007) 1–26

[22] Kelly, F.P.: Reversibility and stochastic networks. Cambridge University Press (2011)

[23] Anderson, D.F., Craciun, G., Kurtz, T.G.: Product-form stationary distributions for deficiency zero chemical reaction networks. B. Math. Biol. 72 (2010) 1947–1970

[24] Anderson, D.F., Cotter, S.L.: Product-form stationary distributions for deficiency zero networks with non-mass action kinetics. B. Math. Biol. 78 (2016) 2390–2407

[25] Innocentini, G.C., Guiziou, S., Bonnet, J., Radulescu, O.: Analytic framework for a stochastic binary biological switch. Phys. Rev. E 94 (2016) 062413

[26] Innocentini, G.C., Forger, M., Radulescu, O., Antoneli, F.: Protein synthesis driven by dynamical stochastic transcription. B. Math. Biol. 78 (2016) 110–131

[27] Zhou, T., Liu, T.: Quantitative analysis of gene expression systems. Quantitative Biology 3 (2015) 168–181

[28] Pendar, H., Platini, T., Kulkarni, R.V.: Exact protein distributions for stochastic models of gene expression using partitioning of Poisson processes. Physical Review E 87 (2013) 042720

[29] Yang, X., Wu, Y., Yuan, Z.: Characteristics of mRNA dynamics in a multi-on model of stochastic transcription with regulation. Chinese J. Phys. 55 (2017) 508–518

[30] Singh, A., Vargas-Garcia, C.A., Karmakar, R.: Stochastic analysis and inference of a two-state genetic promoter model. Proceedings of the American Control Conference, Washington, DC (2013) 4563–4568

[31] Abramowitz, M., Stegun, I.: Handbook of Mathematical Functions with Formulas, Graphs, and Mathematical Tables. National Bureau of Standards, Washington, D.C. (1972)

[32] Cardelli, L., Kwiatkowska, M., Laurenti, L.: Stochastic analysis of chemical reaction networks using linear noise approximation. Biosystems 149 (2016) 26–33

[33] Cinquemani, E.: On observability and reconstruction of promoter activity statistics from reporter protein mean and variance profiles. In: International Workshop on Hybrid Systems Biology, Springer (2016) 147–163

[34] Herath, N., Del Vecchio, D.: Reduced linear noise approximation for biochemical reaction networks with time-scale separation: The stochastic tQSSA+. J. Chem. Phys. 148 (2018) 094108

[35] Bronstein, L., Koeppl, H.: A variational approach to moment-closure approximations for the kinetics of biomolecular reaction networks. J. Chem. Phys. 148 (2018) 014105

[36] Singh, A., Grima, R.: The linear-noise approximation and moment-closure approximations for stochastic chemical kinetics. arXiv preprint arXiv:1711.07383 (2017)

[37] Ghusinga, K.R., Vargas-Garcia, C.A., Lamperski, A., Singh, A.: Exact lower and upper bounds on stationary moments in stochastic biochemical systems. Phys. Biol. 14 (2017) 04LT01

[38] Soltani, M., Vargas-Garcia, C.A., Singh, A.: Conditional moment closure schemes for studying stochastic dynamics of genetic circuits. IEEE Transactions on Biomedical Systems and Circuits 9 (2015) 518–526

[39] Bokes, P., King, J., Wood, A., Loose, M.: Multiscale stochastic modelling of gene expression. J. Math. Biol. 65 (2012) 493–520

[40] Kim, J.K., Josić, K., Bennett, M.R.: The validity of quasi-steady-state approximations in discrete stochastic simulations. Biophys. J. 107 (2014) 783–793

[41] Shahrezaei, V., Swain, P.: Analytical distributions for stochastic gene expression. P. Natl. Acad. Sci. USA 105 (2008) 17256

[42] Popovic, N., Marr, C., Swain, P.S.: A geometric analysis of fast-slow models for stochastic gene expression. J. Math. Biol. 72 (2016) 87–122

[43] Veerman, F., Marr, C., Popović, N.: Time-dependent propagators for stochastic models of gene expression: an analytical method. J. Math. Biol. (2017) 1–52

[44] Platini, T., Jia, T., Kulkarni, R.V.: Regulation by small RNAs via coupled degradation: Mean-field and variational approaches. Phys. Rev. E 84 (2011) 021928

[45] Kumar, N., Jia, T., Zarringhalam, K., Kulkarni, R.V.: Frequency modulation of stochastic gene expression bursts by strongly interacting small RNAs. Phys. Rev. E 94 (2016) 042419

[46] Kurtz, T.G.: The relationship between stochastic and deterministic models for chemical reactions. J. Chem. Phys. 57 (1972) 2976–2978

[47] Kendall, D.: Stochastic processes and population growth. J. Roy. Stat. Soc. B 11 (1949) 230–82

[48] Bokes, P., King, J.R., Wood, A.T., Loose, M.: Exact and approximate distributions of protein and mRNA levels in the low-copy regime of gene expression. J. Math. Biol. 64 (2012) 829–854

[49] Kan, X., Lee, C.H., Othmer, H.G.: A multi-time-scale analysis of chemical reaction networks: II. stochastic systems. J. Math. Biol. 73 (2016) 1081–1129

[50] Lestas, I., Paulsson, J., Ross, N., Vinnicombe, G.: Noise in gene regulatory networks. IEEE T. Circuits-I 53 (2008) 189–200

[51] Feinberg, M.: Lectures on chemical reaction networks. Notes of lectures given at the Mathematics Research Center of the University of Wisconsin in 1979 (1979)

[52] Abou-Jaoudé, W., Thieffry, D., Feret, J.: Formal derivation of qualitative dynamical models from biochemical networks. Biosystems 149 (2016) 70112

[53] Paulsson, J.: Models of stochastic gene expression. Phys. Life Rev. 2 (2005) 157–175

[54] van Kampen, N.: Stochastic Processes in Physics and Chemistry. Elsevier (2006)

[55] Gillespie, D.: A General method for numerically simulating stochastic time evolution of coupled chemical reactions. J. Comput. Phys. 22 (1976) 403–34

[56] Maarleveld, T.R., Olivier, B.G., Bruggeman, F.J.: StochPy: A comprehensive, user-friendly tool for simulating stochastic biological processes. PloS one 8 (2013) e79345

[57] Zhang, F., Wang, D.: The pattern of microRNA binding site distribution. Genes 8 (2017) 296

[58] Mastny, E., Haseltine, E., Rawlings, J.: Two classes of quasi-steady-state model reductions for stochastic kinetics. J. Chem. Phys. 127 (2007) 094106

[59] Srivastava, R., Haseltine, E.L., Mastny, E., Rawlings, J.B.: The stochastic quasi-steady-state assumption: Reducing the model but not the noise. The Journal of chemical physics 134 (2011) 154109

[60] Johnson, N., Kotz, S., Kemp, A.: Univariate Discrete Distributions, 3rd ed. Wiley-Interscience (2005)

[61] Singh, A., Bokes, P.: Consequences of mRNA transport on stochastic variability in protein levels. Biophys. J. 103 (2012) 1087–1096

[62] Ghaemi, R., Del Vecchio, D.: Stochastic analysis of retroactivity in transcriptional networks through singular perturbation. In: American Control Conference (ACC), 2012, IEEE (2012) 2731–2736

[63] Hojcka, M., Bokes, P.: Non-monotonicity of Fano factor in a stochastic model for protein expression with sequesterisation at decoy binding sites. Biomath 6 (2017) 1710217

[64] Soltani, M., Bokes, P., Fox, Z., Singh, A.: Nonspecific transcription factor binding can reduce noise in the expression of downstream proteins. Phys. Biol. 12 (2015) 055002

[65] Bokes, P., Singh, A.: Protein copy number distributions for a self-regulating gene in the presence of decoy binding sites. PloS one 10 (2015) e0120555

